# Deletion of the AMPylase mFICD alters cytokine secretion and affects cognitive plasticity *in vivo*

**DOI:** 10.1101/2021.04.16.440208

**Authors:** Nicholas McCaul, Corey M Porter, Anouk Becker, Chih-Hang Antony Tang, Charlotte Wijne, Bhaskar Chatterjee, Djenet Bousbaine, Angelina M Bilate, Chih-Chi Andrew Hu, Hidde L Ploegh, Matthias C Truttmann

**Affiliations:** Program in Cellular and Molecular Medicine, Boston Children’s Hospital, Boston, MA, 02115, USA; Harvard Medical School, Boston, MA, 02115, USA; Department of Molecular & Integrative Physiology, University of Michigan, Ann Arbor, MI, 48109, USA; Geriatrics Center, University of Michigan, Ann Arbor, MI, 48109, USA; Center for Translational Research in Hematologic Malignancies, Houston Methodist Cancer Center, Houston Methodist Research Institute, Houston, TX 77030, USA

**Keywords:** post-translational modification, AMPylation, FICD, HYPE, BiP, UPR

## Abstract

Fic domain-containing AMP transferases (fic AMPylases) are conserved enzymes that catalyze the covalent transfer of AMP to proteins. This post-translational modification regulates the function of several proteins, including the ER-resident chaperone Grp78/BiP. Here we introduce a mFICD AMPylase knock-out mouse model to study fic AMPylase function in vertebrates. We find that mFICD deficiency is well-tolerated in unstressed mice. We show that mFICD-deficient mouse embryonic fibroblasts are depleted of AMPylated proteins. mFICD deletion alters protein synthesis and secretion in splenocytes, including that of IgM and IL-1β, without affecting the unfolded protein response. Finally, we demonstrate that older mFICD^-/-^ mice show improved cognitive plasticity. Together, our results suggest a role for mFICD in adaptive immunity and neuronal plasticity *in vivo*.

## Introduction

The post-translational regulation of protein function is a fundamental concept in biology. To manage protein activity, dedicated enzymes attach specific chemical modifications to individual proteins, the presence of which affect the behavior and activity of the modified proteins. These modifications, called post-translational modifications (PTMs), govern essential biological processes. They are implicated in cancer, neurodegeneration and cardiovascular diseases, among others.

The covalent addition of an AMP moiety to the side chain of exposed threonine and serine residues has emerged as a new paradigm to control the activity of the essential ER-resident chaperone BiP. This process, AMPylation, is catalyzed by metazoan AMP transferases (AMPylases) that contain a *filamentation induced by c-AMP* (fic) domain. Fic domain-containing AMPylases (fic AMPylases) are highly conserved and are present in a single copy in most metazoans, including *Caenorhabditis elegans* (FIC-1), *Drosophila melanogaster* (dfic), *Mus musculus* (mFICD) and *Homo sapiens* (FICD)^1-3^.

Metazoan fic AMPylases are bi-functional: using a single active site, these enzymes catalyze both the transfer of AMP to surface-exposed threonine and serine hydroxyl groups, and the removal of AMP groups from modified residues (deAMPylation)^4-8^. The switch between AMPylation and deAMPylation is proposed to involve enzyme dimerization, the exchange of Mg^2+^ with Ca^2+^ ions in the active site, and the subsequent switch from an open to a closed conformation^4-9^. The latter is stabilized by interactions between an inhibitory glutamate and a nearby arginine residue, which aligns an inhibitory α-helix such that the catalytic core preferentially binds AMP over ATP, catalyzing deAMPylation^6^. If the interactions between these residues are prevented or resolved, fic AMPylases adopt an open conformation that favors Mg^2+^ and ATP recruitment to the active site, enabling AMPylation of target proteins^10^. Thus, replacing the critical inhibitory glutamate residue with a glycine (FICD(E234G)) converts the enzyme to a constitutively-active AMPylase^10-12^.

AMPylation of the ER-resident HSP70 protein, BiP, on T518 locks this chaperone in an ATP- and HSP40-bound “primed” conformation, rendering it unable to support the (re)folding of client proteins^7,13^. Upon BiP deAMPylation, ATP is hydrolyzed and the ADP-bound form of BiP is again able to engage with client proteins^6,14^. The consequences of BiP S365/T366 AMPylation remain controversial and may either inhibit^4,12,15^ or enhance^16,17^ BiP activity. In addition to BiP, fic AMPylases also modify a wide range of non-ER proteins^18-27^. Indeed, fic AMPylases are also present in the nuclear envelope and the cytoplasm^11,20,28^.

Changes in cellular AMPylation levels affect cellular fitness and organismal survival: Over-expression of constitutively active fic AMPylases is toxic and kills human^17,29,30^ and yeast cells^20^, as well as worm (*Caenorhabditis elegans*) embryos^19^ and flies (*Drosophila melanogaster*)^4^. In contrast, fic AMPylase deficiency is well-tolerated in unstressed human cells but impairs the activation of the unfolded protein response (UPR) under stress^17^ and reduces neuronal differentiation^25^. Further, Fic-1 deficient worms show enhanced sensitivity to the presence of aggregation-prone poly-glutamine repeat proteins in neurons^19^. Perhaps the most significant *in vivo* fic AMPylase knock-out (KO) phenotype is found in dfic deficient flies, which show significant defects in visual signaling and suffer from light-induced blindness caused by BiP deregulation^5,31^. Despite the emerging role of fic AMPylases in proteostasis, our understanding of how these enzymes affect mammalian physiology is lacking.

Here we describe the generation and characterization of a mFICD deficient mouse strain. mFICD^-/-^ mice are viable and are not visually impaired. We further show that mFICD deletion alters IgM synthesis and perturbs IL-1β secretion. Finally, we demonstrate that mFICD^-/-^ mice show improved cognitive function as they age. Together, our results support a role for mFICD function in adaptive immunity and neuronal plasticity in vertebrates.

## Results

### mFICD deficient mice are viable and fertile

To investigate the role of mFICD-mediated protein AMPylation *in vivo*, we attempted to generate both mFICD-deficient as well as constitutively active mFICD(E234G)-expressing transgenic mouse strains using CRISPR/Cas9 technology. We used a sgRNA that targets a site adjacent to the coding sequence of the regulatory motif (TVAIEG) (Fig. S1A) and a double stranded repair template to introduce the E234G substitution. The injection of approximately 80 blastocysts resulted in more than 20 independent animals carrying insertions or deletions in the mFICD gene that often resulted in frame shifts. Notably, not a single animal carrying the constitutively active mFICD(E234G) mutation was recovered. Parallel injections using identical experimental conditions but targeting different genes efficiently produced transgenic knock-in strains^32^. These results suggest that embryonic expression of constitutively active mFICD(E234G), particularly in the absence of a wild-type mFICD copy, may be lethal.

For this study, we backcrossed a mouse strain carrying a deletion in mFICD, which introduced a frame shift resulting in a premature stop codon (Fig. S1B-C). To characterize the mFICD^-/-^ animals, we assessed 6 months-old female control and mFICD^-/-^ animals using the SHIPRA method. SHIPRA is a rapid, comprehensive screening approach, which provides a qualitative behavioral and functional profile for each animal^33^. We found that mFICD^-/-^ mice performed similarly to control animals for all 14 assessed features (Fig. 1A; feature by feature results in table S1). We found no significant differences in body weight (Fig. 1B), rotarod performance (Fig 1C) and lifespan (Fig. S1D) between control and mFICD^-/-^ animals. Together, these results establish that mFICD deficiency is well tolerated by mice under normal growth conditions.

**Figure 1:**
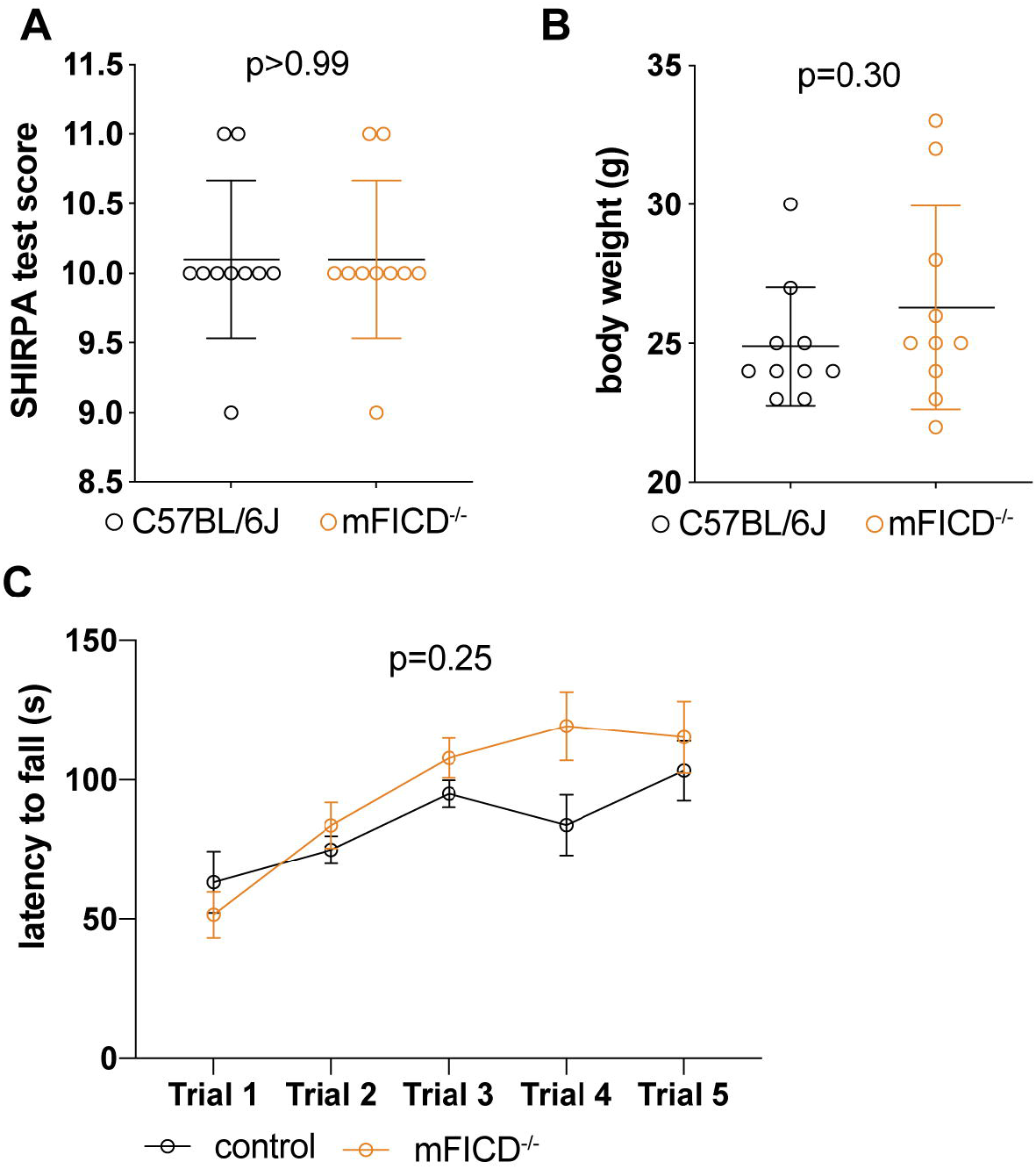
mFICD^-/-^ mice are viable and do not show obvious phenotypes. SHIPRA test scores, body weight (B) latency to fall from rotarod (C) of 6 months-old wild-type and mFICD^-/-^ mice. n=10/cohort. Error bars represent SD. Statistical significance (P values) were calculated using two-way ANOVA for repeated measures with Geisser-Greenhouse correction (C) or unpaired t-tests (A and B).

### mFICD is required for AMPylation of BiP, EEF-1A and HSC70 in mouse embryonic fibroblasts (MEFs)

AMPylation in vertebrates is conferred by at least two evolutionarily unrelated enzymes: FICD and the mitochondrial pseudokinase SelO^34^. To define how mFICD deficiency alters the vertebrate AMPylome, we supplemented mFICD^-/-^ and control mouse embryonic fibroblast (MEF) lysates with 8-azido-ATP. Following AMPylation, a click reaction was used to install a PEG-biotin handle on the modified proteins. We then recovered AMPylated proteins with streptavidin-coated beads and identified AMPylated proteins by mass spectrometry (Fig. 2A). Comparing results from treated MEFs, mFICD^-/-^ KO MEFs and a cell-free control, we identified 108 proteins that were AMPylated only in wild-type MEFs (Table S2). Among these proteins were several known FICD targets, including BiP (HSPA5), HSC70 (HSPA8), and translation elongation factor EEF-1A (Fig. 2A, S2A). In contrast, only two proteins (transketolase (TKT); Protein disulfide isomerase (P4HB)) were identified in both wild-type and mFICD^-/-^ lysates. Gene ontology analysis showed that AMPylated proteins were significantly enriched in metabolic, protein (re-)folding and stress response processes (Fig. S2B). Together, these results confirm that FICD is required for the AMPylation-mediated regulation of BiP, HSC70, EEF-1A and other proteins.

**Figure 2:**
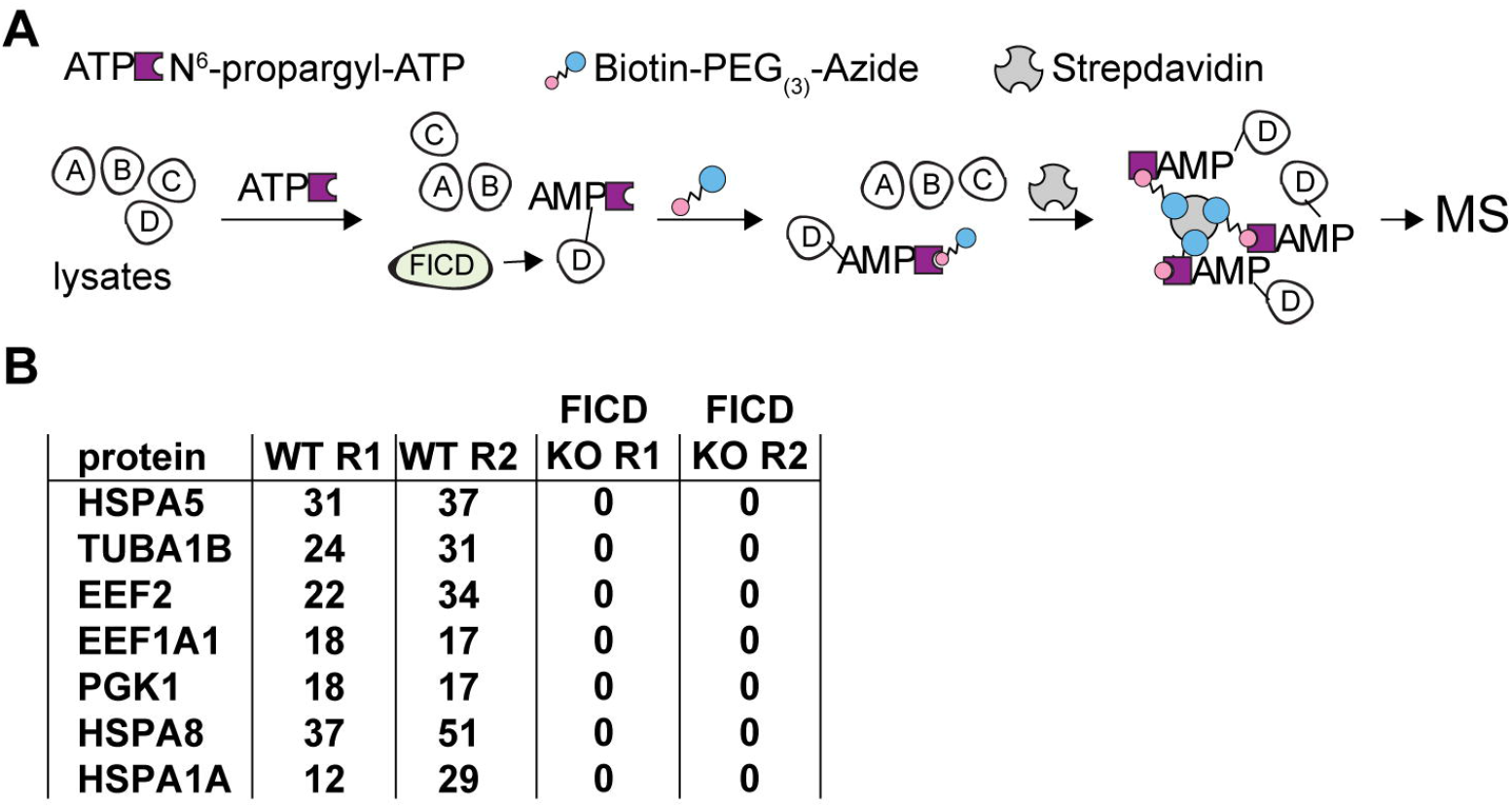
Deletion of mFICD almost completely abrogates protein AMPylation. (A) schematic representation of experimental setup. (B) Table showing comparative enrichment of indicated proteins in wild-type and MEF KO cells.

### B and T cell development is unaffected by mFICD deletion

Having established that WT and mFICD^-/-^ mice are phenotypically similar in overall physiological and morphological terms, and given mFICD’s role in the UPR, an important pathway in B and T cell development^35^ we asked whether their humoral immune system was affected by mFICD deficiency. Using flow cytometry, we examined the distribution of B and T cells in the spleen. We found no difference between WT and mFICD^-/-^ mice in either the distribution of B and T cells (Fig. 3A and S3A), nor in any T cell (Fig. 3B and S3B) or B cell (Fig. 3C and S3C) subsets present in the spleen. B cell development in the bone marrow (Fig. 3D and S3D) and T cell development in the thymus (Fig. 3E and S3E) were normal in mFICD^-/-^ mice.

**Figure 3:**
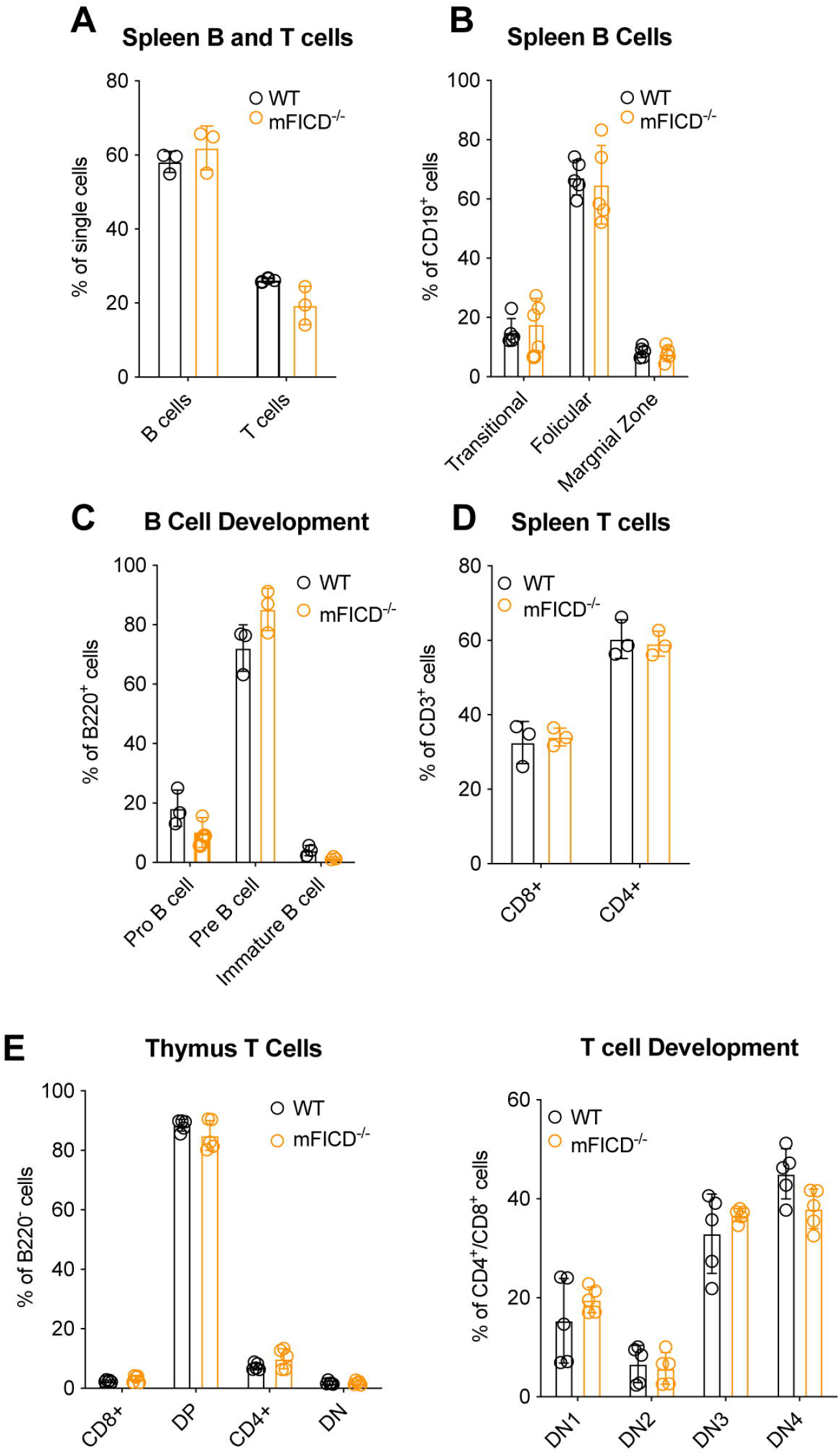
B and T cell development is normal in mFICD^-/-^ mice. Splenocytes from age-matched WT and mFICD^-/-^ mice were stained with antibodies against immune-cell markers and analyzed by flow cytometry to determine B and T cell populations (A) and relevant B cell (B) and T cell (D) subsets. (D) Flow cytometry was performed as in (A) on cells isolated from bone marrow of WT and mFICD^-/-^ mice to follow B cell development. (E) Flow cytometry was performed as in (A) on cells isolated from thymus of WT and mFICD^-/-^ mice to observe T cell populations (E) and follow T cell development (F). Each circle represents 1 data from 1 mouse, n ≥ 3 mice per experiment.

### Deletion of mFICD perturbs protein secretion in splenocytes

BiP, an essential molecular chaperone, discovered as an immunoglobulin-binding protein and regulated by FICD, is required for antibody assembly and maturation. We thus examined the impact of mFICD deficiency on protein secretion. Splenocytes were isolated from WT and mFICD^-/-^ mice and stimulated with lipopolysaccharide (LPS), heparan sulfate (HS) or thapsigargin to induce cytokine secretion, which was then assayed by ELISA (Fig. 4A-C). While mFICD deletion significantly reduced LPS-induced IL-6 secretion (Fig. 4A), it had no effect on secretion of TNFα (Fig. 4B). We also observed a significant difference in IL-1β production between WT and mFICD^-/-^ mice (Fig. 4C). This is striking because unlike IL-6 and TNFα, which traffic through the ER, IL-1β folds in the cytoplasm and is secreted by a non-classical pathway. Immunoblots showed that intracellular levels of the IgM heavy chain (µ) were elevated in mFICD^-/-^ B cells, while the levels of other proteins that fold and traffic through the ER were unchanged (Fig. S4A).

**Figure 4:**
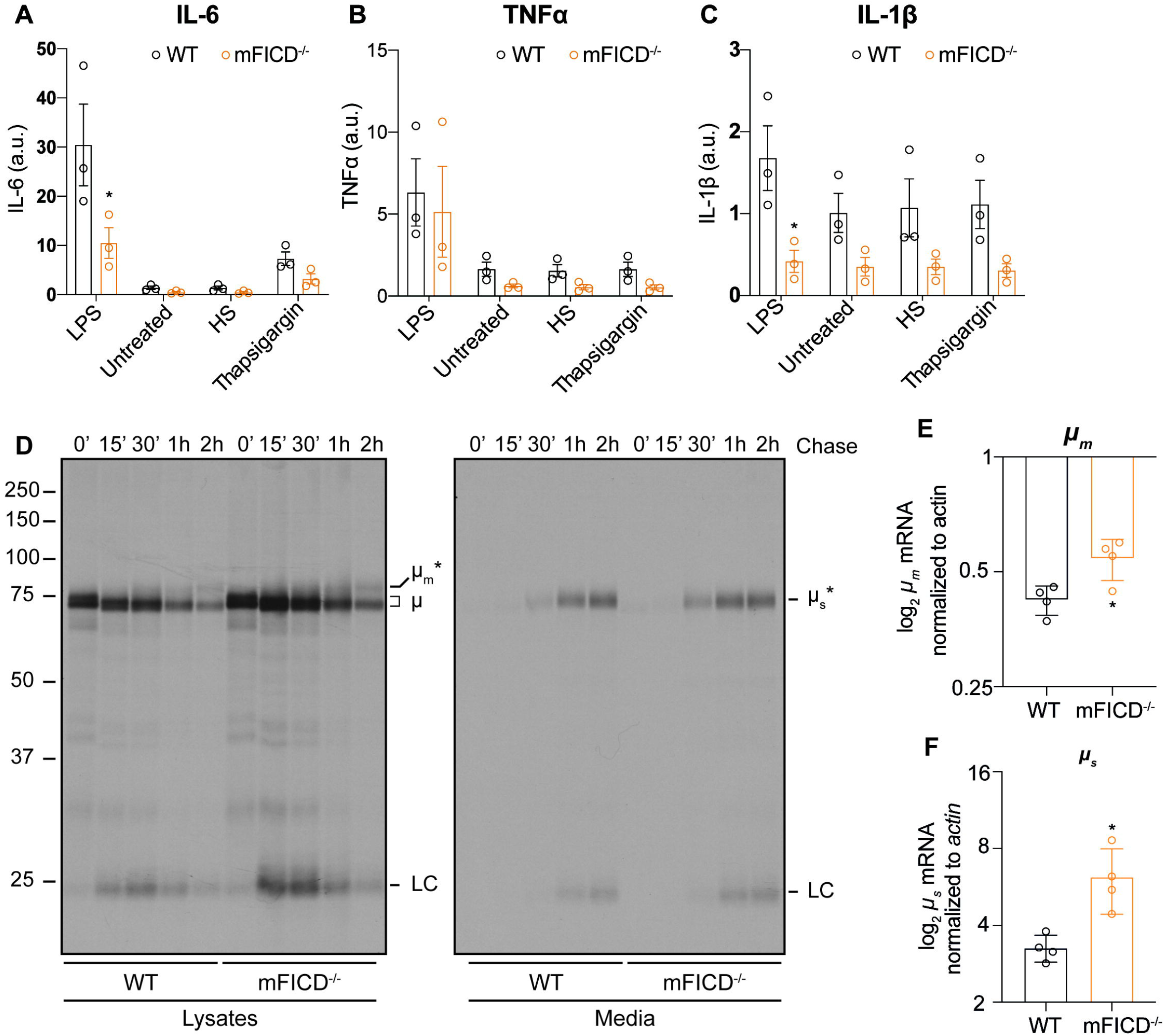
Protein secretion in mFICD^-/-^ splenocytes and B cells is perturbed. (A-C) Splenocytes were isolated from WT and mFICD^-/-^ mice and incubated with lipopolysaccharide (LPS), heparan sulfate (HS) or thapsigargin (Tg) and monitored for secretion of IL-6 (A), IL-1β (B) and TNFα (C). (D) 3-day LPS stimulated B cells were pulse labeled with 35S methionine and cysteine and chased for the indicated times. Media samples and detergent lysates were immunoprecipitated with an anti-µ polyclonal antibody and then analyzed by SDS-PAGE and autoradiography. Gels are representative of 5 independent experiments. qPCR was performed on RNA purified from 3-day LPS activated B cells as in (D) using primers against membrane-bound (E) or soluble (F) µ. Primers against actin were used for normalization. *=p<0.05.

To further explore the consequences of mFICD deletion on protein folding and secretion, we focused on immunoglobulins because of their requirement for BiP activity in the course of folding and assembly^36^. Naïve B cells were purified from spleens, activated using LPS and cultured for 3 days to allow differentiation into IgM-secreting plasmablasts. We then followed IgM folding by pulse-chase analysis. Briefly, plasmablasts were labeled with ^35^S-methionine/cysteine for 15 minutes and chased in the absence of radioactive label for various times to follow protein maturation. IgM was immunoprecipitated from detergent lysates and media samples and analyzed by SDS-PAGE followed by autoradiography (Fig. 3D). mFICD^-/-^ mice showed increased levels of both the soluble µ heavy chain (µ_s_) and the membrane-bound B cell receptor (µ_m_) in detergent lysates. This was accompanied by increased levels of both µ_s_ and µ_m_ mRNA (Fig. 4E and F). Similar amounts of IgM were recovered from the media for both WT and mFICD^-/-^ samples, suggesting that the increased levels of µ in the mFICD^-/-^ cells did not pass ER quality control for secretion. These observations were consistent across the 5 pulse-chase experiments performed. We did not observe differences in the synthesis or glycosylation of other ER-folding proteins (Fig. S4B).

### mFICD deletion does not impair induction of the unfolded protein response

AMPylation of BiP exerts fine control over the level of active BiP present in the ER^7,12,13,17^. To examine how abrogation of such control affects the folding capacity and stress tolerance of cells, we examined the physiological UPR induced during B cell activation. As before, we isolated B cells from total splenocytes and stimulated them with LPS. Immunoblots showed no change in the levels of UPR sensors IRE1α or PERK, both before and after stimulation (Fig. 5A). There was also no change in the activation of either receptor as monitored by XBP1 splicing or eif2α phosphorylation, or in the expression of downstream targets BiP and Grp94 (Fig. 5A). We further verified this by examining downstream targets of the UPR by qPCR (Fig. 5B). We observed no changes in the expression levels between WT and mFICD^-/-^ samples for any of the genes that are downstream targets of the IRE1α, PERK and ATF6α pathways.

**Figure 5:**
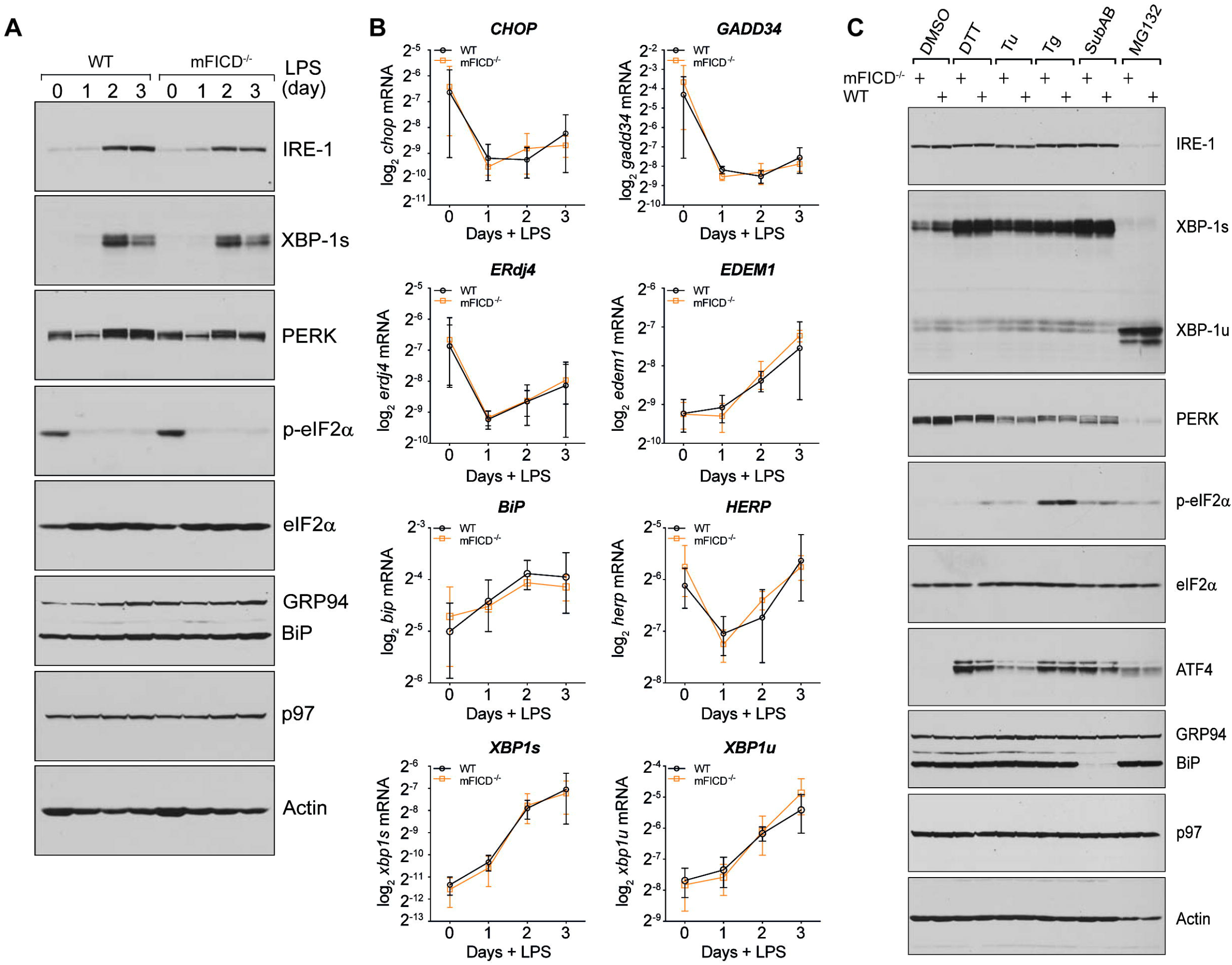
mFICD deletion does not alter UPR in plasmablasts. (A) Naïve B cells purified from spleens of WT and mFICD^-/-^ mice were stimulated with LPS for a course of 3 days to allow for differentiation into plasmablats and samples were collected each day for protein analysis via immunoblot. Post-nuclear supernatants were analyzed by SDS-PAGE and immunoblotting with the indicated antibodies. (B) Samples collected as in (A) were lysed in TriZol and used for RNA purification and analysis by qPCR using primers directed against the indicated genes. Actin was used as reference gene. (C) Naïve B cells similarly purified as in (A) were stimulated with LPS for 2 days and subsequently incubated with DMSO, DTT (5 mM; 3 h), Tunicamycin (Tu; 10 µg/mL; 8 h), Thapsigargin (Tg; 2.5 µM; 8 h), Subtilase cytotoxin (SubAB; 100 ng/mL; 8 h) or MG132 (50 µM; 8 h). These treated plasmablasts cells were then lysed and subjected to analysis by immunoblot as in (A).

The physiological UPR induced upon B cell activation is anticipatory of enhanced antibody production and differs from a UPR induced by an accumulation of misfolded proteins. To ascertain whether mFICD^-/-^ cells responded differently to the latter type of UPR, we incubated B cells with several chemical initiators of the UPR. As with the physiological UPR from B cell activation, we found no significant difference in any of the UPR receptors or downstream targets assayed by immunoblot (Fig. 5C).

### mFICD^-/-^ mice are not visually impaired

In *Drosophila melanogaster*, the mFICD ortholog dfic regulates reversible photoreceptor degeneration, which is critical for visual neurotransmission and adaptation to constant light exposure^5,31^. We thus tested whether mFICD^-/-^ mice show signs of visual impairment. As part of the SHIRPA test (see table S1), we assessed visual placing and the pupillary light reflex, which was normal in all tested 6 months-old control as well as mFICD^-/-^ animals. To confirm these results, we performed optometry tests, in which mice were presented a rotating grating, a condition that elicits head movements in mice with intact eyesight^37^. mFICD deficiency did not impact the animal’s response to the moving grating early (6 months old) or late (18 months old) in life (Fig. 6A-B). These results suggested that mFICD deficiency does not affect vision in mice.

**Figure 6:**
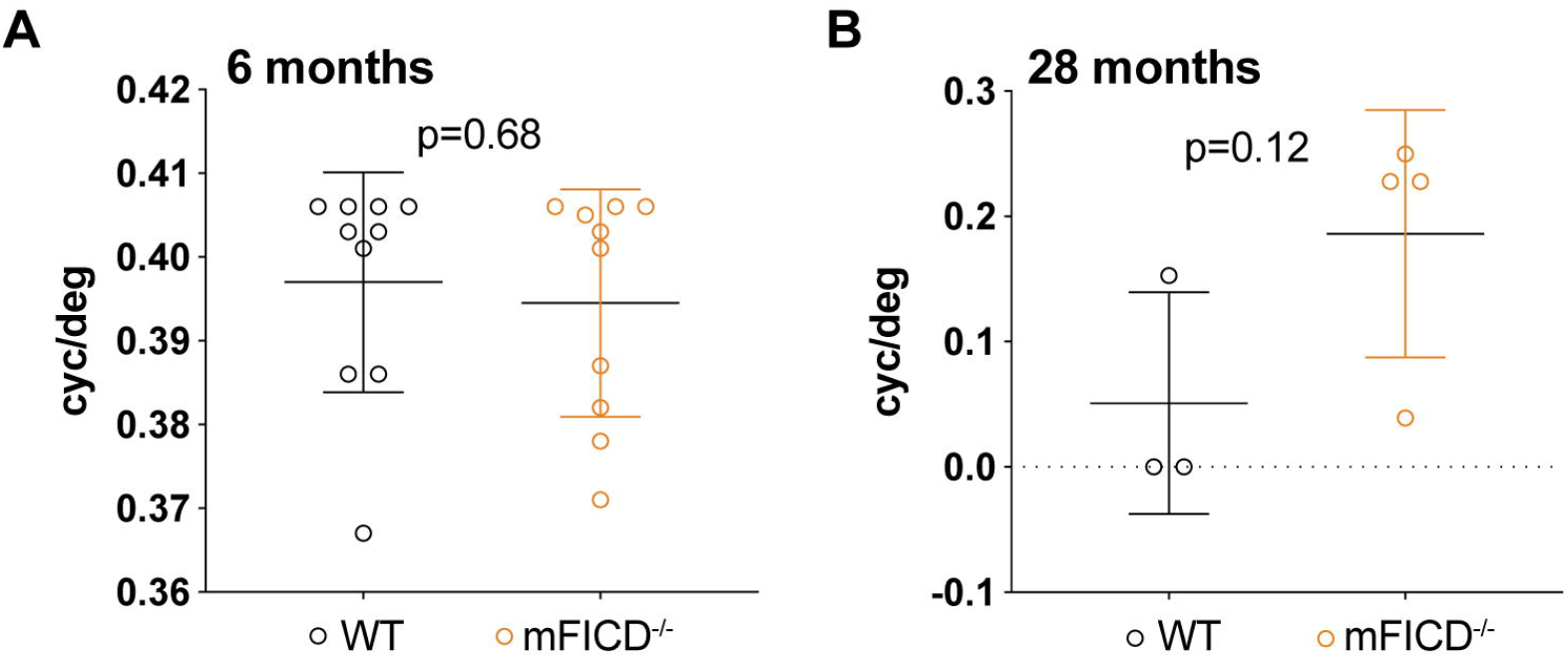
mFICD^-/-^ mice do not visual impairments. Optometry-based assessments of visual perception of 6-months (A) and 28 months old wild-type and mFICD^-/-^ mice. Statistical significance (P values) were calculated using unpaired t-tests.

### mFICD deficiency does not impair learning, cognition and memory

FICD activity modulates neurogenesis and neuronal differentiation in human cerebral organoids^25^. To determine if mFICD^-/-^ animals may have defects in learning, cognition and memory, we tested control and mFICD^-/-^ animals at 6 and 18 months of age in Morris water maze tests^38^. These tests are designed to assess impairments in visual short-term and long-term memory and visual-spatial abilities by observing and recording escape latency, distance moved and velocity^38^. There was no significant difference in visual nonspatial short-term learning comparing wild-type and mFICD^-/-^ animals at 6 months of age (Fig. 7A-B). However, 18-month old mFICD^-/-^ mice significantly decreased escape latency between trial 1 and 2 while the performance of wild-type mice did not (Fig 7C-D). Next, we assessed learning and memory capacity of mFICD^-/-^ and wild-type mice at 6 and 18 months of age using a submerged platform setup. Both control and mFICD^-/-^ mice performed equally well (Fig. 7E, S5A) phase. Upon removal of the platform, wild-type and mFICD^-/-^ showed similar recall behaviors, spending the most time in the maze quadrant that contained the platform in the preceding learning phase (Fig. 7F-G). Swimming speed was similar between control and mFICD^-/-^ cohorts (Fig. S5B), suggesting that mFICD deficiency does not affect aging-dependent decline in rough muscle function. Together, these results indicate that cerebellar function, learning and memory are not affected by mFICD loss. Finally, to test for cognitive flexibility, we reintroduced the platform to a new position (quadrant) in the water maze. We found that both control and mFICD^-/-^ mice at 6 and 18 months of age adapted to the new situation and memorized the new position of the platform equally well (Fig. 7H).

**Figure 7:**
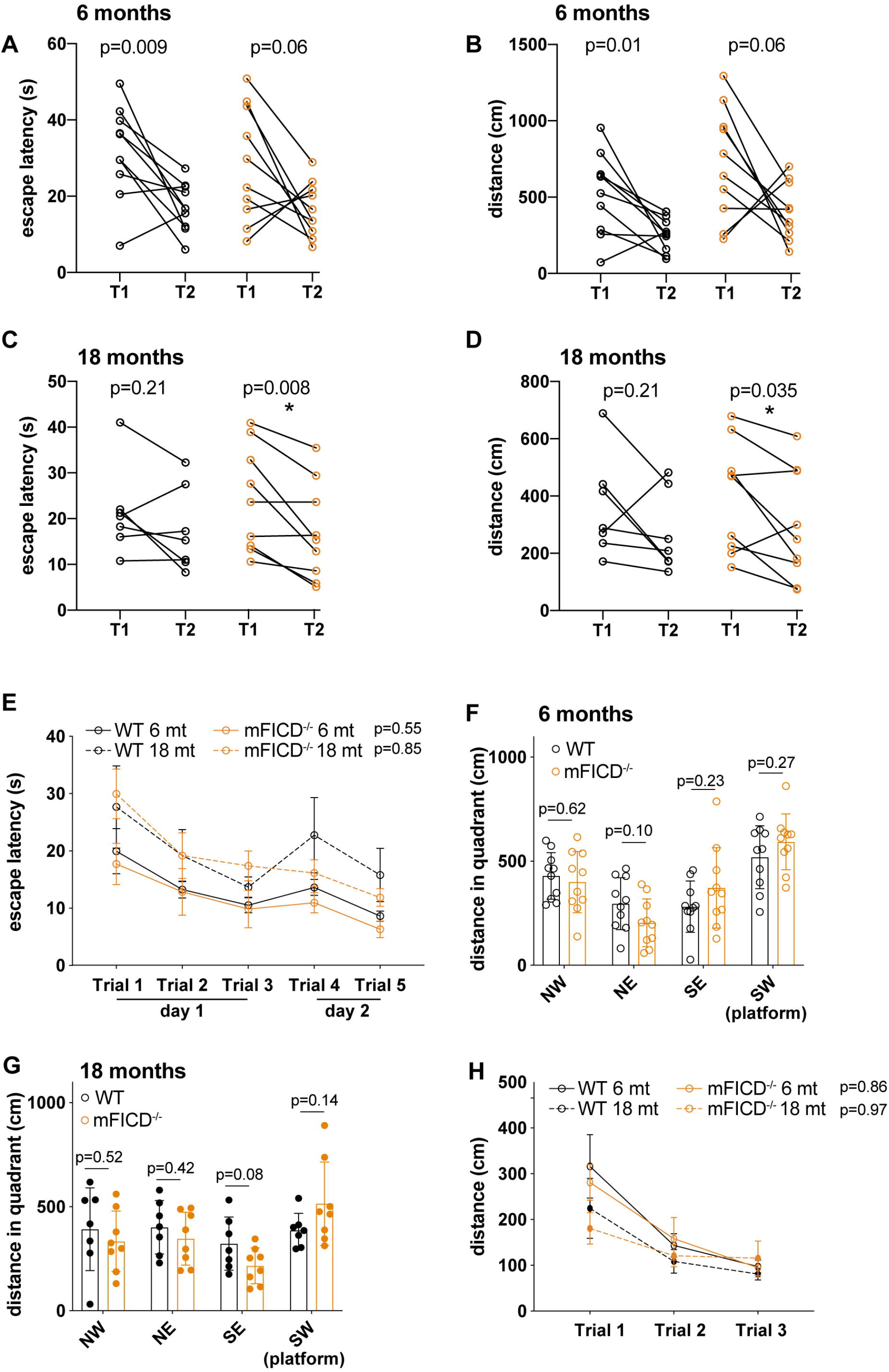
mFICD^-/-^ mice show no signs of cognitive deficits. Morris water maze tests to assess visual and spatial learning and memory. Escape latency (A, C) and distance (B, D) of 6 months and 18 month old mice in initial learning test using visible platform. (E) escape latency of 6 and 18 months old mice in subsequent learning phase using a submerged platform. (F-G) distance in quadrant (recall of platform) of 6 months and 18 months old mice. (H) distance to platform upon moving the platform to new position (cognitive flexibility measure). Statistical significance (P values) were calculated using two-way ANOVA for repeated measures with Geisser-Greenhouse correction (B and D) or unpaired t-tests (A and C).

## Discussion

Protein AMPylation in metazoans is increasingly recognized as a PTM that regulates the function of the ER-resident chaperone BiP as well as that of other proteins ^15,21^. While the consequences of hyper- and hypo-AMPylation are well established in tissue culture systems and invertebrate models, the impact of fic AMPylase deficiency on mammalian physiology is unknown. Here we describe a functional mFICD knockout mouse. We have examined the physiological, neurological and molecular consequences of mFICD deletion and identify points where WT and mFICD^-/-^ mice differ. Overall, our phenotypic and pathologic assessments of mFICD^-/-^ mice did not identify major debilitating abnormalities or dysfunctions. These findings are in accordance with previous work in human tissue culture and *C. elegans in vivo* models for FIC-1 deficiency, both of which showed that the absence of FICD/FIC-1 is well-tolerated in unstressed cells and animals^5,7,11,12,17^. In contrast with work done in dfic-deficient flies, which suffer from deficits in visual perception and light-induced blindness^5,31^, we did not observe differences or defects in vision in mFICD^-/-^ mice. Possible explanations for these divergent observations include the presence of compensatory/alternative mechanisms to regulate proteins AMPylated by mFICD^-/-^ in mice and/or differences in target protein profile between dfic and mFICD. Further work is required to decipher the role of mFICD in visual perception in molecular detail.

On a molecular level, mFICD deficiency depletes unstressed cells of most AMPylated proteins. In accordance with previous studies, we find evidence that mFICD AMPylates both ER-resident as well as cytoplasmic proteins^11,19,20,25-29,31,39^. The mechanisms that underlie mFICD’s ability to target topologically distinct compartments remain to be defined. AMPylation of BiP by mFICD regulates the levels of active BiP to allow a fine-tuned response to ER stress^7^. We show that in B cells, abrogation of mFICD activity resulted in higher levels of µ synthesis in the ER but secretion of IgM is similar for WT and mFICD^-/-^ B cells. BiP binds to the CH1 domain of immunoglobulins to prevent aggregation and promote folding, only releasing the heavy chain molecule once a correctly folded light chain is available for association. Increased levels of intracellular µ could be due to a deficit in light chain levels or possibly due to strengthened interaction between the CH1 domain and BiP due to increased BiP activity.

We further discovered that deletion of mFICD compromised secretion of IL-1β. Pro-IL-1β folds in the cytoplasm where it is cleaved by caspase-1 to produce the active cytokine, which is then released from the cell via non-classical secretion through gasdermin-D pores^40^. IL-1β can enter both the type I and type III non-conventional protein secretion pathways^41^. It is therefore possible that AMPylation plays a role in the regulation of non-classical secretion. Of note, the bacterial Hsp70 (DnaK) has also been implicated in non-classical secretion^42^.

mFICD^-/-^ mice do not show signs of cognitive impairment; rather, they had a tendency to perform somewhat better in most assays than wild-type control animals, a trend we also observed in motor assays. We attribute this to the presumably continuous de-repression of BiP and the dysregulation of other AMPylated proteins such as HSC70 and EEF-1A in the absence of mFICD. Future work on mFICD-dependent and indepdent protein AMPylation in the presence of aggregation-prone proteins or in response to stressors that cause protein unfolding should define the role of this PTM in neuronal health and aging.

In conclusion, our work shows that mFICD deficiency is tolerated in the absence of stress but can impair BiP-dependent (antibody folding and maturation) and - independent (IL-1β secretion) processes. Future studies focusing on the effects of protein AMPylation in the ER and beyond should provide additional insights into proteostasis and protein folding.

## Materials and Methods

### Generation of FICD^-/-^ mouse

All animal procedures were performed according to NIH guidelines and approved by the Committee on Animal Care at MIT and the Institutional Animal Care and Use Committee (IACUC) at Boston Children’s Hospital. CRISPR/Cas9-mediated genome editing was performed exactly as described in Maruyama *et al*^32^. We targeted a genomic location using a sgRNA (ccacacggtggccatcga<u>ggg</u>) close to the sequence encoding for the regulatory FICD motif (TVAIEG), which resulted in multiple transgenic animal containing 1-300 bp long deletion or insertions. Mice were out-crossed 6 times with C57BL/6J animals to eliminate putative off-target mutation introduced during CAS9-based genome editing.

### Identification of AMPylated proteins

Mass spectrometry-based identification of AMPylated proteins was performed using a chemical reporter setup as previously described^43,44^. Briefly, we supplemented total lysates of mFICD^-/-^ and wild-type MEFs with N6-propargyl-ATP and incubated the lysates at room temperature for 1 hour. This allowed endogenous mFICD to utilize N^6^-propargyl-ATP as nucleotide substrate. We then supplemented the reaction with biotin-(PEG)_3_-azide to covalently couple a biotin handle to the AMP-propargyl groups now found on AMPylated proteins. AMPylated (biotinylated) proteins were recovered using Streptavidin-modified agarose beads, eluted and analyzed by LC/MS/MS.

### Behavioral test

We examined 10 female C57BL/6J (wild type) and 10 female mFICD^-/-^ mice setup in cages of four, each containing two control and two mFICD^-/-^ animals. Each mouse received a unique paw tattoo to enable ID tracking throughout the experiment. All project members involved in behavioral testing of animals were blinded and unaware of cage compositions.

### SHRIPA test

SHIRPA testing was performed as described previously^33,45^.

### Immunoblotting

B cells or LPS-stimulated plasmablasts were lysed in RIPA buffer (10 mM Tris-HCl, pH 7.4; 150 mM NaCl; 1% NP-40; 0.5% sodium deoxycholate; 0.1% SDS; 1 mM EDTA) supplemented with protease inhibitors (Roche) and phosphatase inhibitors. Protein concentrations were determined using BCA assays (Pierce). Protein samples were boiled in SDS-PAGE sample buffer (62.5 mM Tris-HCl, pH 6.8; 2% SDS; 10% glycerol; 0.1% bromophenol blue) with β-ME, analyzed by SDS-PAGE, and transferred to nitrocellulose membranes, which were subsequently blocked in 5% non-fat milk (wt/vol in PBS), and immunoblotted with indicated primary antibodies and appropriate horseradish peroxidase-conjugated secondary antibodies (Southern Biotech). Primary antibodies to IRE-1 (Cell Signaling Technology), XBP-1 (Cell Signaling Technology), PERK (Santa Cruz), phospho-eIF2α (Ser51; Cell Signaling Technology), eIF2α (Cell Signaling Technology), ATF4 (Cell Signaling Technology), GRP94/BiP (anti-KDEL; Enzo Life Sciences), p97 (Fitzgerald), actin (Sigma-Aldrich), µ heavy chain (Southern Biotech), and κ light chian (Southern Biotech) were obtained commercially. Polyclonal antibodies against mouse class I MHC heavy chain, class II MHC α, β or invariant (li) chains, Igβ and STING were generated in rabbits. Immunoblots were developed using Western Lightning Chemiluminescence Reagent (Perkin-Elmer).

### Morris water maze tests

Mice were placed in a circular pool (137-cm diameter) filled with water maintained at room-temperature. The tank was divided into quadrants each marked on the wall of the tank with a different visual cue for spatial orientation. Swim time and path length were recorded by automated video tracking (Ethovision XT 11.5, Noldus Information Technology, The Netherlands). A black curtain surrounded the tank to prevent cues in the room aiding the spatial performance. On Day 1 (visible platform), mice were given 90 s to swim to a platform elevated 1 cm above the water and marked with a flag. The position of the platform was kept constant, but the starting quadrant varied. Four starts (each from one of the four quadrants) were administered in one trial, and two visual trials were administered during day 1. On Days 2 and 3 (learning) the platform was moved to the opposite quadrant from the visible position, submerged 1 cm below the water line and the flag was removed. Mice were given 90 s to find the hidden platform and left on the platform for 5 s to orientate themselves. Mice that did not find the platform within 90 s were guided to it by the experimenter. Each mouse completed three learning trials on day 2 (Trials 1 to 3), and two learning trials on day 3 (Trials 4 and 5), with each trial comprising four different start positions. On Day 4 (recall), the platform was removed entirely from the tank, the mice placed in the quadrant opposite the quadrant that had previously contained the platform (e.g. north east, NE), and given 60 s to swim. Time spent and path length traversed in each of the four quadrants was recorded. On Day 5 (recall), the platform was placed back into the tank 1cm below the water in a quadrant different from the visible and learning trials and mice were tested across 3 trials (4 starts of 90 s each from different locations).

### Rotarod tests

On day 1, mice were placed on a rotarod (Economex, Columbus Instruments, USA) set to revolve at 4 rpm for 5 continuous minutes. This training phase allows mice to acclimate to the movement of the rotarod. The following day, mice are placed on the rotarod revolving at 4 rpm for a 10 second acclimation before accelerating at 0.1 rpm/sec. These trials were repeated 4 times with a break of at least 5 minutes between repeats. Latency to fall was recorded as read out.

### Optometer tests

We used a protocol adapted from Prusky *et al*.^37^, which employs an optomotor device (CerebralMechanics, Canada) to measure visual acuity. The device consisted of 4 computer monitors arranged in a square with a lid on top to enclose the mouse within. A computer program was used to project on the monitors a virtual cylinder in 3-D coordinate space. Visual stimuli were drawn on the walls of the cylinder, and from the perspective of the platform, each monitor appeared as a window on a surrounding 3-D world. The software also controlled the speed of rotation and geometry of the cylinder and the spatial frequency and contrast of the stimuli. A red crosshair in the video frame indicated the center of the cylinder rotation. Mice were placed one at a time on the platform inside the device, the lid of the box was closed, and the animals were allowed to move freely and as the mouse moved about the platform, the experimenter followed the mouse’s head with the red crosshair. When a grating perceptible to the mouse was projected on the cylinder wall and the cylinder was rotated (12 deg/sec), the mouse normally stopped moving its body and would begin to track the grating with reflexive head movements in concert with the rotation. An experimenter assessed whether the animals tracked the cylinder by monitoring in the video window the image of the cylinder, the animal, and the crosshair simultaneously. If the mouse’s head tracked the cylinder rotation, it was judged that the animal could see the grating. Using a staircase procedure, the mouse was tested systematically against increasing spatial frequencies of the grating until the animal no longer responded. The threshold was then calculated as the highest spatial frequency that the mouse responded to.

### Cytokine measurements

Total splenocytes were isolated from C57BL/6 and mFICD^-/-^ mouse spleens and then treated for 3 days with 20 μg/ml LPS (Sigma), 100 µg/ml heparan sulfate or 2.5 μM thapsigargin (Enzo Life Sciences). Supernatants were harvested after 72 h and TNFα, Il-6 and IL-1β levels were quantified by ELISA (Biolegend kits).

### Cell culture

Nave B lymphocytes were purified from mouse spleens by negative selection using CD43 (Ly48) magnetic beads (Miltenyi Biotec) according to the manufacturer’s instructions.. Naïve B cells were cultured in RPMI 1640 media (Gibco) supplemented with 10% heat-inactivated FBS, 2 mM L-glutamine, 100 U/mL penicillin G sodium, 100 μg/mL streptomycin sulfate, 1 mM sodium pyruvate, 0.1 mM nonessential amino acids, and 0.1 mM β-mercaptoethanol (β-ME).

### Flow Cytometry

WT and mFICD^-/-^ mice aged 6-8 weeks were euthanized by CO_2_ asphyxiation followed by cervical dislocation. Spleen, bone marrow and thymus tissues were extracted and homogenized in PBE buffer (PBS + 0.5% BSA and 1 mM EDTA). Red blood cells were lysed using ACK lysis buffer (Gibco) and cells were resuspended in PBE for surface staining with the following antibodies (clone;source) for 30 minutes at 4°C: B220 PerCP-Cy5.5, CD43 APC, CD19 Pacific Blue, IgM FITC, IgD APC-Cy7, CD21 PerCP-Cy5.5, CD23 FITC, IgM APC, CD44 FITC, CD25 PE-Cy7, CD4 APC, CD8α Pacific Blue (53-6.7;BioLegend), CD3 PerCP-Cy5.5 (145-2C11;BD), B220 APC-Cy7. All samples were blocked using Fc-Block (BD Bioscience). Acquisition of B and T cell populations was performed on an LSRFortessa cytometer (BD) instrument and analyzed with the FlowJo software package (Tree-Star).

### Pulse chase labelling and immunoprecipitation

Suspension pulse chase was performed as previously described^46^. Briefly, LPS-stimulated plasmablasts were starved for methionine and cysteine for 60 minutes and pulse labeled for 15 minutes with 55 µCi/1×10^6^ cells of Express ^35^S protein labeling mix (Perkin Elmer). Samples were then diluted 10x with cell culture media containing 5mM unlabeled methionine and cysteine. At the end of each chase time, a sample was removed and diluted 1:1 into ice-cold PBS containing 20 mM N-ethyl maleimide (Sigma). Cells were collected by centrifugation at 1250 xg for 5 minutes and washed twice with ice-cold PBS. Cells were lysed in RIPA buffer supplemented with protease inhibitors (Roche) and phosphatase inhibitors.. IgM was immunoprecipitated from lysates or media samples with a goat anti-mouse IgM antibody (Southern Biotech). Immunoprecipitates were washed with lysis buffer and eluted from beads with reducing SDS-PAGE sample buffer and analyzed by SDS-PAGE. Gels were then dried and exposed to BioMax MR films (Carestream).

### RNA extraction, cDNA synthesis and qPCR

LPS-activated plasmablasts on each day of activation were lysed using TriZol (Invitrogen) and RNA was isolated using RNA extraction kit (Zymo). RNA was checked for integrity and equivalent amounts of RNA were converted into cDNA using the Maxima H minus cDNA Synthesis master mix (Invitrogen). Samples for qPCR were prepared with primers against target genes (Table 1) and mixed with SSO Advanced Universal SYBR Green Supermix (BioRad) in 96-well plates and analyzed on a CFX96 qPCR machine (BioRad). Analysis was performed using standard Δct methods using actin as a reference gene.

### Statistical analysis

Data is graphed as means +/-SEM; where feasible, individual datapoints are shown. Depending on the experiment, we used unpaired/paired two-tailed t tests, Turkey’s multiple comparison’s test, 2-way ANOVA with multiple comparison test as well as Kaplan-Meier tests to evaluate survival data. Individual p values are displayed in each figure.

## Acknowledgements

We like to thank the Truttmann, Ploegh and Hu laboratories for feedback on the manuscript. The Animal Behavior and Physiology Core at Boston Children’s Hospital is acknowledged for performing the behavioral tests (IDDRC 1U54HD090255). This work was supported by grants NIH/NCI R01CA163910 to C.C.H. and support from AARG and MICHR UL1TR002240 to M.C.T.

---

**Figure S1:**
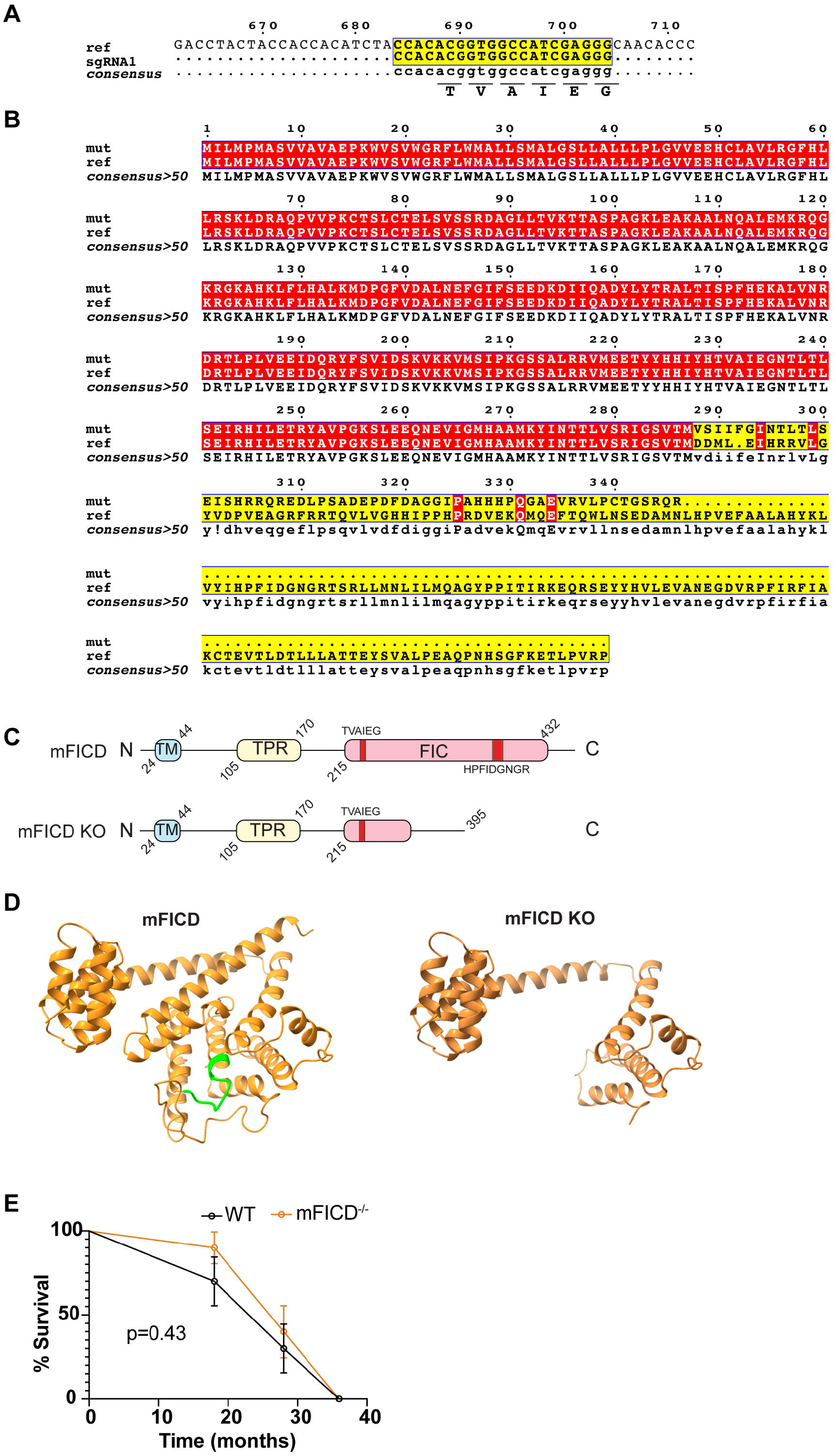
generation of mFICD-/-mice using CRISPR/Cas9. (A) schematic of experimental strategy. sgRNA was chosen based on its proximity to the regulatory TVAIEG motif. (B) In silico translation of mFICD-/-allele. (C) Schematic of wild-type and mFICD KO-encoded mFICD. TM: transmembrane domain; TPR: tetratricopeptide repeat domain; FIC: filamentation induced by cyclic AMP (catalytic) domain. (D) Survival of mice after 6, 16 and 28 months. Statistical significance (P=0.99) was calculated using Log rank (Mantel Cox) test.

**Figure S2:**
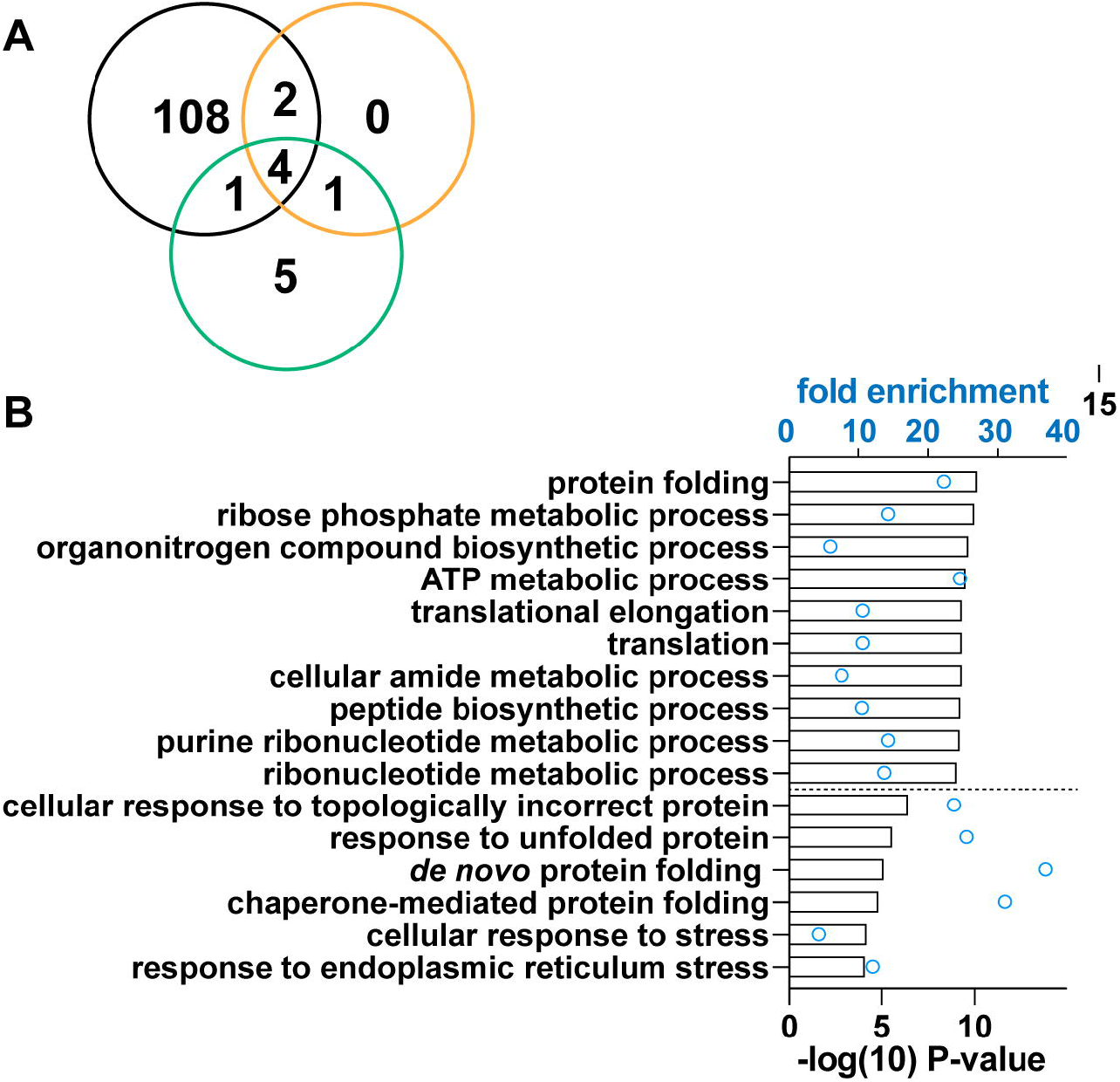
Gene ontology (GO) analysis of AMPylome. (A) Venn diagram showing the number of proteins AMPylated in wild-type (black circle), mFICD knockout (orange circle) MEFs and a cell-free control (turquoise circle). (B) GO enrichment analysis was performed comparing AMPylated proteins to reference proteome.

**Figure S3:**
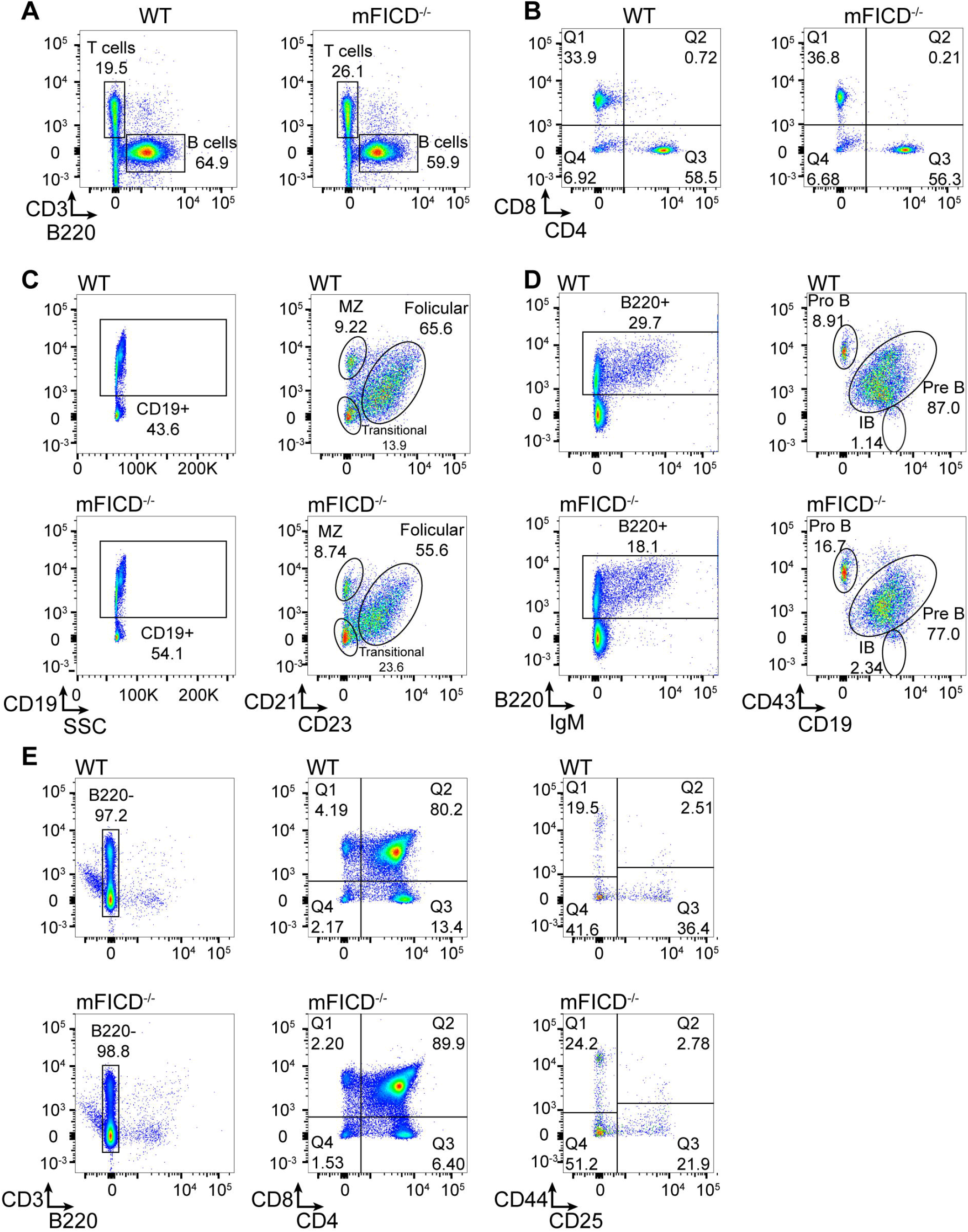
B and T cell development is normal in mFICD^-/-^ mice. Splenocytes from age-matched WT and mFICD^-/-^ mice were stained with antibodies against immune-cell markers and analyzed by flow cytometry to determine B and T cell populations (A), T cell subsets (B) and B cell subsets (C). (D) Flow cytometry was performed as in (A) on cells isolated from the bone marrow of WT and mFICD^-/-^ mice to follow B cell development. (E) Flow cytometry was performed as in (A) on cells isolated from thymus of WT and mFICD^-/-^ mice to follow T cell populations. (F) CD4/CD8 populations from (E) were further examined to follow T cell development. FACS plots are representative of multiple mice in each group.

**Figure S4.**
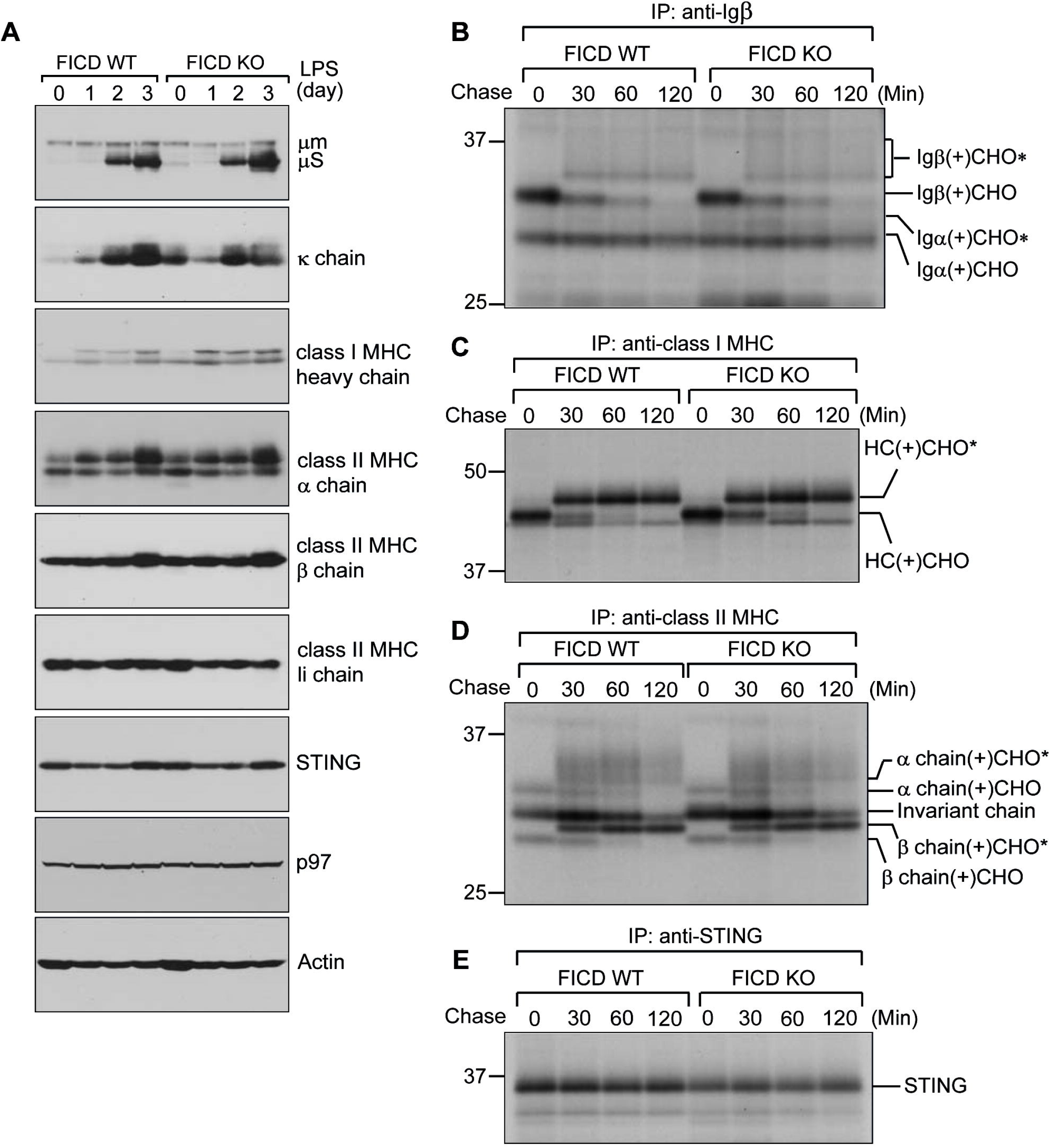
Immune receptor levels and glycoprotein trafficking are normal in mFICD-/-plasmablasts. (A) Splenocytes were isolated from WT and mFICD-/-mice and incubated with lipopolysaccharide (LPS) for the indicated times. Post-nuclear supernatants were analyzed by SDS-PAGE, transferred to nitrocellulose membranes and analyzed by immunoblot for the indicated targets. (B-E) 3 day LPS-activated plasmablasts were pulsed and chased as in Figure 3D. Detergent lysates were immunoprecipitated with antibodies against Igβ (B), Class I MHC (C), Class II MHC (D), and STING (E). Immunoprecipitates were analyzed by SDS-PAGE and autoradiography.

**Figure S5.**
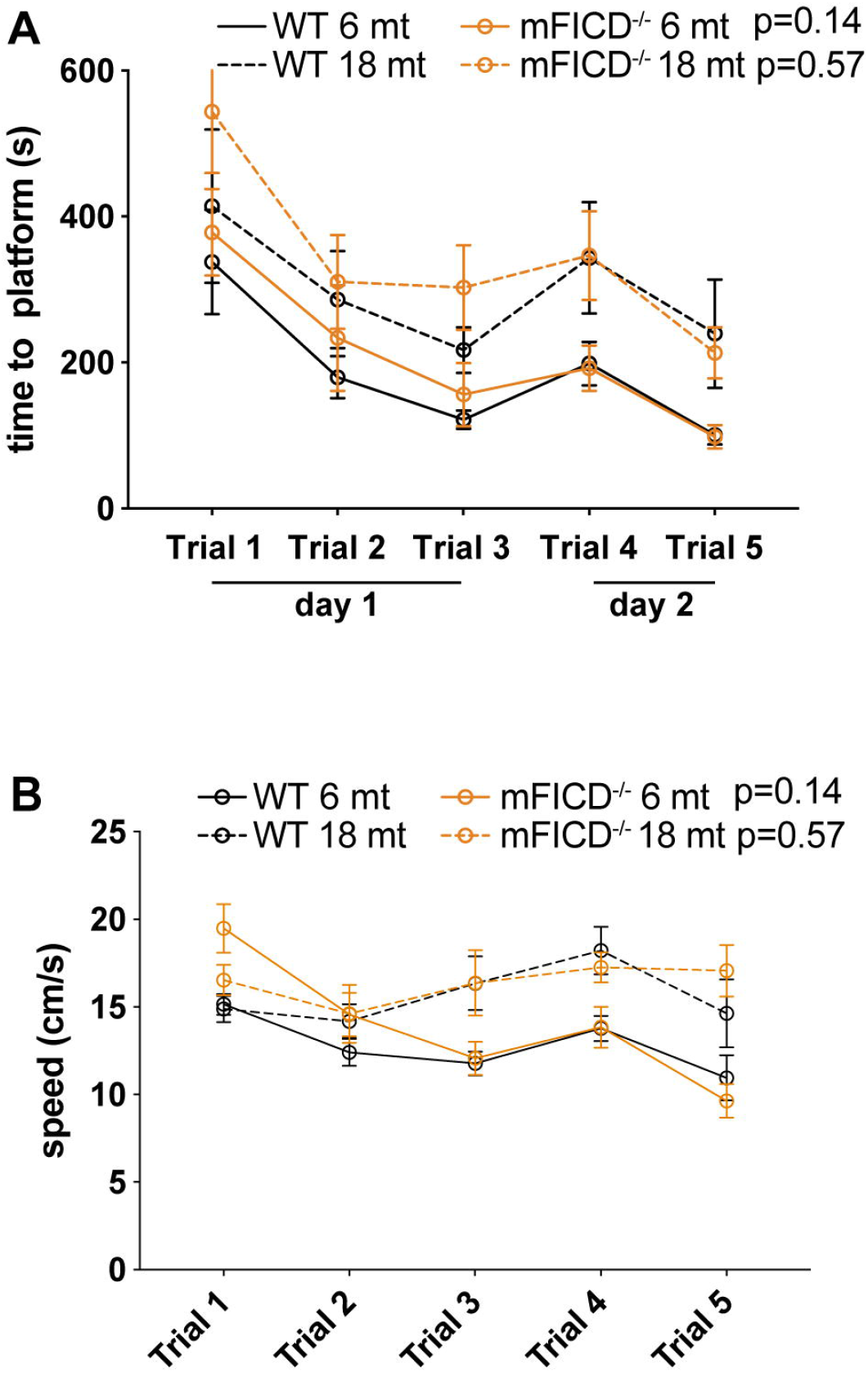
mFICD^-/-^ mice show no signs of cognitive deficits. Morris water maze tests to assess visual and spatial learning and memory. (A) time to platform of 6 and 18 months old mice. (B) Average swimming speed of 6 and 18 months old animals. Statistical significance (P values) were calculated using two-way ANOVA for repeated measures with Geisser-Greenhouse correction (C).

## Notes

### Competing Interest Statement

The authors have declared no competing interest.

